# FUNGI DIVERSITY IN THE RHIZOSPHERE OF *Aspilia pruliseta* Schweif. ext Schweif IN THE SEMI-ARID EASTERN KENYA

**DOI:** 10.1101/2020.10.23.351908

**Authors:** James Peter Muchoka, Daniel Njiru Mugendi, Paul Nthakanio Njiruh, Paul Kamau Mbugua, Ezekiel Mugendi Njeru, Amanuel Menghs Ghilamicael, Mariciano Iguna Mutiga

## Abstract

Semi-arid eastern Kenya is a fragile ecosystem with continuous cultivation of dryland pulses and grains. Farmers use artificial fertilizers most of which are deleterious to the environment. Previous studies have shown that soil microbes in the rhizosphere could be used to sustainably enhance levels of soil mineral nutrients and soil health. However, few studies have examined fungal diversity in the rhizosphere of wild and native *Aspilia pruliseta* shrub. In this study, amplicons of Internal Transcribed Spacer (ITS) region on Total Community DNA using Illumina sequencing were used to explore the fungal community composition within the rhizosphere. Operational taxonomic units (OTUs) were analyzed using QIIME 1.8.0, taxonomy assigned via BLASTn against SILVA 119 database. Hierarchical clustering was done using R programming software. 72,093, 50,539 and 43,506 sequence reads were obtained from samples MC1_a_, MC2_a_ and MC3_a_ respectively representing rhizosphere depth 0-20 cm, 21-40 cm and 41-60 cm. A total of 373 OTUs were realized at 3% genetic distance. Taxonomic analysis revealed that the genera *Glomus* was most prevalent in all soil depths with 85.60 % of the OTUs in depth 0-20 cm, 69.04 % in depth 21-40 cm and 48.45 % in depth 41-60 cm. The results revealed high levels of obligate arbuscular mycorrhiza fungi that if commercially cultured could enhance phosphates uptake in crops.

## Introduction

Microorganisms were used in early civilizations for agricultural and industrial processes long before they were well known and documented. Recent advancements in understanding about the genetics, physiology, and biochemistry of fungi, has led to the exploitation of fungi for different purposes in agriculture and industrial products of economic importance. Application of chemical fertilizers to crop plants negatively affects human health and environments (1). Beneficial plant-microbes interactions in the rhizosphere are determinants of plant health and soil fertility (2). Recent studies have focused on identification of alternative methods to enhance plant productivity and protect the soil. This can only be possible with thorough knowledge and information of microbes along the plants’ rhizosphere and fungal microbes is one category, that with this understanding, could provide the link and alleviate challenges posed by poor land productivity.

Arbuscular mycorrhizal fungi (AMF) are below ground symbiotic associations between plant roots and fungi (3). AMF can improve plant growth under low fertility conditions, improve water balance of the plants and help plants to establish in new areas (4). These beneficial effects imply that the plant community structure and productivity in ecosystems are influenced significantly by the AM fungal diversity in the soil (5). However, the genetic diversity of fungi and other microbes in the plants’ rhizosphere is not yet fully comprehended (1).

Fungi dominate in low pH or slightly acidic soils where soils tend to be undisturbed (6). A good undisturbed ecological niche would be a plants’ rhizosphere. Over 80% of vascular or non-vascular terrestrial plants form symbiotic mycorrhizae fungi relationships along the rhizosphere by forming hyphae networks (7). Through mycorrhizae the plant obtains mainly phosphate and other minerals, such as zinc and copper, from the soil. The fungus obtains nutrients, such as sugars, from the plant root. Mycorrhiza fungi interdependence with roots of the higher plants is not only beneficial to the host plant but also play a role in the aggradative process of soil structure formation (8). Furthermore, a well diversed rhizosphere fungal community is thought to play a role in the suppression of pathogens (9) and (10). Knowledge of the structure and diversity of the fungal community in the rhizosphere will lead to a better understanding of pathogen-antagonist interactions (10).

Most past plant rhizosphere mycological studies have concentrated on cultivated crops (11) (12) and (13). This has left a gap in understanding fungal diversity preexisting before planting field crops. In this study, illumina sequencing molecular method was used to identify fungal species in the rhizosphere of *Aspilia pruliseta* Schweif. *Aspilia pruliseta* is a flowering plant in the asteraceae family and was hypothesized to grow in coexistence with the mycorrhiza fungi. It is hypothesized that this complex association leads to availability of phosphorus in the soils in usable forms to plants. The herbaceous plant is common and grows naturally in the open woodlands and grasslands in western, southern, central and eastern Africa. Farmers in central eastern Kenya have reported good crop yields, particularly cereals, in farms previously growing *Aspilia pruliseta*.

## MATERIALS AND METHODS

### Study sites

This study was carried out in the semi-arid eastern Kenya at Gakurungu, Tunyai and Kanyuambora with coordinates 00^0^21’00” S, 37^0^28’30” E, 00^0^10’00” S, 37^0^50’00” E and 00^0^12’00” S, 37^0^51’00” E respectively. Elevation for all the studied sites was below 1000 m above sea level. Rainfall in the selected areas was scanty and below 700 mm/year with two distinct, but unreliable wet seasons in the months of March to May and October to December. Dry spells were more prolonged with temperature mean of 26^0^C. Natural trees and shrubs found in the area were *Aspilia pruliseta*, *Cassia sp, Euphorbia sp, Acasia sp* and *Balanites aegyptiaca*. *Cenchrus ciliaris* and *Hyperrhenia rufa* grasses were interspersed in trees and shrubs.

### Measurements of physico-chemical information on soil rhizosphere depths in the study sites

Physico-chemical analysis of the rhizosphere soil was done for factors that would influence fungal populations and distribution (Table 1a, b, c). Latitude and longitude of the sampling sites were taken using global positioning system (GARMIN eTrex 20). Soil pH for each rhizosphere depth was taken with a portable pH meter (Oakton pH 110, Eutech Instruments Pty. Ltd) and confirmed with indicator strips (Merck, range 5-10). *In situ* soil temperature was taken using an electrical chemical analyzer (Jenway – 3405). 10 g of the composite soil from every rhizosphere depth studied was analyzed in the laboratory for mycorrhiza fungi determination using (14) protocol. The same soil sample was analysed for soil phosphorus in ppm using (15) protocol. Soil moisture was determined using (16) method while soil nitrogen content was determined using (17) method. The samples was also analysed for soil phosphates using (18) protocol and organic matter content using the method by (19).

**Table 1 (a).**
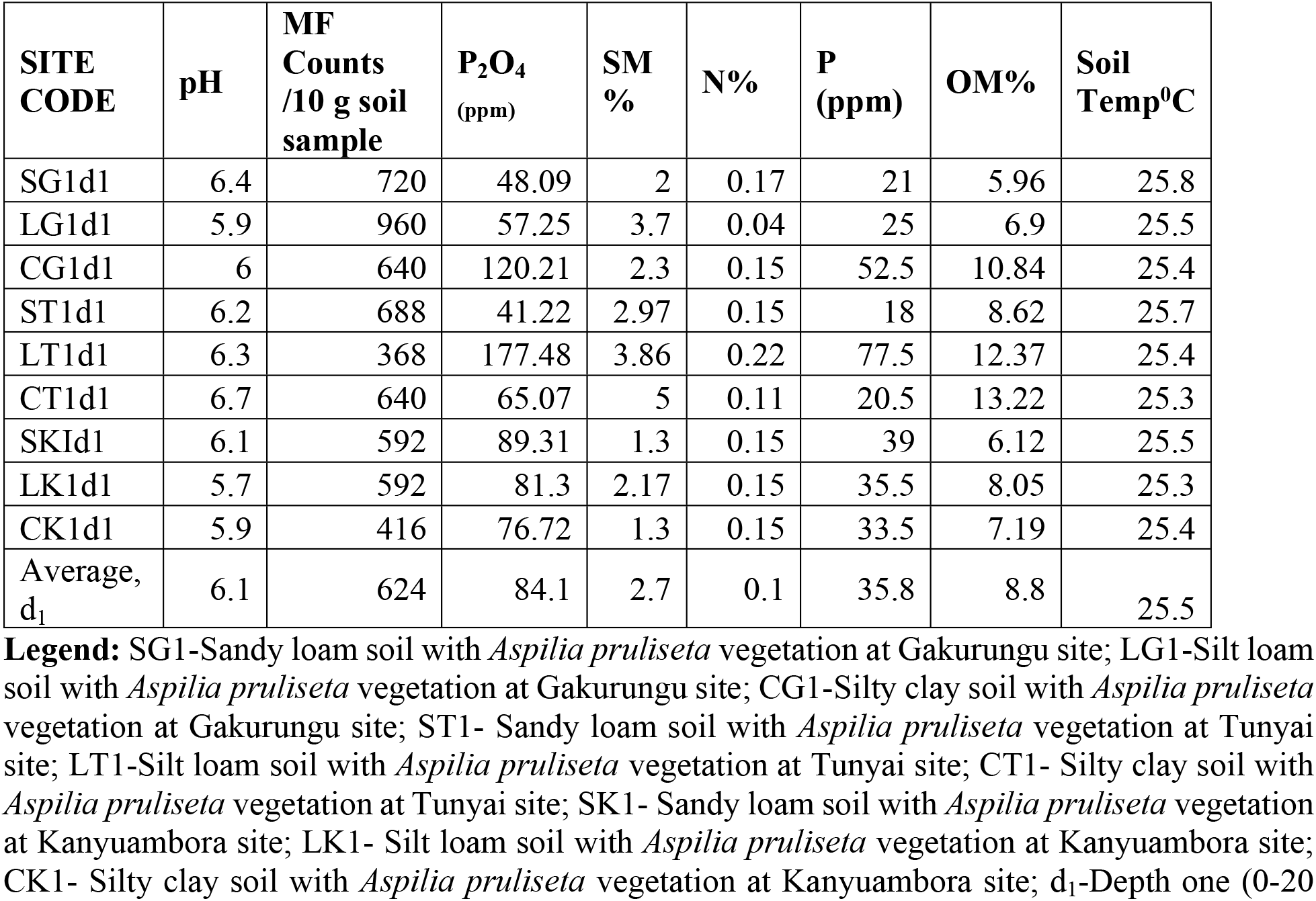
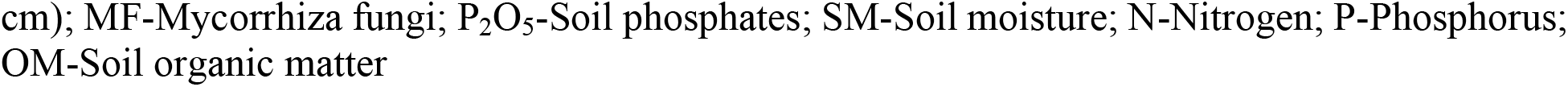
Soil rhizosphere depth_1_ (0-20 cm) physico-chemical parameters in sampling sites

| SITE CODE | pH | MF Counts /10 g soil sample | P<sub>2</sub>O<sub>4</sub> (ppm) | SM % | N% | P (ppm) | OM% | Soil Temp<sup>0</sup>C |
| --- | --- | --- | --- | --- | --- | --- | --- | --- |
| SG1d1 | 6.4 | 720 | 48.09 | 2 | 0.17 | 21 | 5.96 | 25.8 |
| LG1d1 | 5.9 | 960 | 57.25 | 3.7 | 0.04 | 25 | 6.9 | 25.5 |
| CG1d1 | 6 | 640 | 120.21 | 2.3 | 0.15 | 52.5 | 10.84 | 25.4 |
| ST1d1 | 6.2 | 688 | 41.22 | 2.97 | 0.15 | 18 | 8.62 | 25.7 |
| LT1d1 | 6.3 | 368 | 177.48 | 3.86 | 0.22 | 77.5 | 12.37 | 25.4 |
| CT1d1 | 6.7 | 640 | 65.07 | 5 | 0.11 | 20.5 | 13.22 | 25.3 |
| SK1d1 | 6.1 | 592 | 89.31 | 1.3 | 0.15 | 39 | 6.12 | 25.5 |
| LK1d1 | 5.7 | 592 | 81.3 | 2.17 | 0.15 | 35.5 | 8.05 | 25.3 |
| CK1d1 | 5.9 | 416 | 76.72 | 1.3 | 0.15 | 33.5 | 7.19 | 25.4 |
| Average, d<sub>1</sub> | 6.1 | 624 | 84.1 | 2.7 | 0.1 | 35.8 | 8.8 | 25.5 |

**Legend:** SG1-Sandy loam soil with *Aspilia pruliseta* vegetation at Gakurungu site; LG1-Silt loam soil with *Aspilia pruliseta* vegetation at Gakurungu site; CG1-Silty clay soil with *Aspilia pruliseta* vegetation at Gakurungu site; ST1- Sandy loam soil with *Aspilia pruliseta* vegetation at Tunyai site; LT1-Silt loam soil with *Aspilia pruliseta* vegetation at Tunyai site; CT1- Silty clay soil with *Aspilia pruliseta* vegetation at Tunyai site; SK1- Sandy loam soil with *Aspilia pruliseta* vegetation at Kanyuambora site; LK1- Silt loam soil with *Aspilia pruliseta* vegetation at Kanyuambora site; CK1- Silty clay soil with *Aspilia pruliseta* vegetation at Kanyuambora site; d<sub>1</sub>-Depth one (0-20
cm); MF-Mycorrhiza fungi; P<sub>2</sub>O<sub>5</sub>-Soil phosphates; SM-Soil moisture; N-Nitrogen; P-Phosphorus; OM-Soil organic matter

**Table 1(b).** Soil rhizosphere depth_2_ (21-40 cm) physico-chemical parameters in sampling sites

| SITE CODE | pH | MF Counts /10 g soil sample | P<sub>2</sub>O<sub>4</sub> (ppm) | SM % | N% | P (ppm) | OM% | Soil Temp0C |
| --- | --- | --- | --- | --- | --- | --- | --- | --- |
| SG1d2 | 6.7 | 160 | 25.19 | 1.8 | 0.2 | 11 | 6.15 | 25.6 |
| LG1d2 | 5.7 | 176 | 65.27 | 3.48 | 0.1 | 28.5 | 3.34 | 25.4 |
| CG1d2 | 5 | 640 | 25.19 | 2.09 | 0.12 | 11 | 9.89 | 25.3 |
| ST1d2 | 5.7 | 336 | 80.15 | 3.18 | 0.13 | 35 | 10.36 | 25.5 |
| LT1d2 | 5.8 | 280 | 64.12 | 3.66 | 0.14 | 28 | 12.91 | 25.3 |
| CT1d2 | 7.4 | 296 | 80.15 | 5.41 | 0.07 | 35 | 12.53 | 25.1 |
| SK1d2 | 6.1 | 336 | 61.83 | 2.09 | 0.18 | 27 | 9.42 | 25.4 |
| LK1d2 | 4.7 | 320 | 208.39 | 2.78 | 0.01 | 91 | 7.66 | 25.3 |
| CK1d2 | 5.8 | 384 | 46.95 | 1.39 | 0.11 | 20.5 | 9.84 | 25.3 |
| Average d2 | 5.9 | 325.3 | 73 | 2.8 | 0.1 | 31.9 | 9.1 | 25.4 |

**Legend:** SG1-Sandy loam soil with *Aspilia pruliseta* vegetation at Gakurungu site; LG1-Silt loam soil with *Aspilia pruliseta* vegetation at Gakurungu site; CG1-Silty clay soil with *Aspilia pruliseta* vegetation at Gakurungu site; ST1- Sandy loam soil with *Aspilia pruliseta* vegetation at Tunyai site; LT1-Silt loam soil with *Aspilia pruliseta* vegetation at Tunyai site; CT1- Silty clay soil with *Aspilia pruliseta* vegetation at Tunyai site; SK1- Sandy loam soil with *Aspilia pruliseta* vegetation at Kanyuambora site; LK1- Silt loam soil with *Aspilia pruliseta* vegetation at Kanyuambora site; CK1- Silty clay soil with *Aspilia pruliseta* vegetation at Kanyuambora site; d<sub>2</sub>-Depth two (21-40 cm); MF-Mycorrhiza fungi; P<sub>2</sub>O<sub>5</sub>-Soil phosphates; SM-Soil moisture; N-Nitrogen; P-Phosphorus; OM-Soil organic matter

**Table 1(c).** Soil rhizosphere depth_2_ (41-60 cm) physico-chemical parameters in sampling sites

| SITE CODE | pH | MF Counts /10 g soil sample | P<sub>2</sub>O<sub>4</sub> (ppm) | SM % | N% | P (ppm) | OM% | Soil Temp0C |
| --- | --- | --- | --- | --- | --- | --- | --- | --- |
| SG1d3 | 6.6 | 224 | 53.82 | 1.85 | 0.24 | 23.5 | 6.51 | 25.5 |
| LG1d3 | 6 | 208 | 50.38 | 2.57 | 0.14 | 22 | 5.24 | 25.3 |
| CG1d3 | 5 | 384 | 246.75 | 1.69 | 0.12 | 10.15 | 7.22 | 25.1 |
| ST1d3 | 5.6 | 144 | 73.28 | 2.89 | 0.09 | 32 | 6.99 | 25.4 |
| LT1d3 | 5.9 | 200 | 95.04 | 4.76 | 0.04 | 41.5 | 15.82 | 25.2 |
| CT1d3 | 8 | 160 | 44.66 | 5.25 | 0.05 | 19.5 | 12.58 | 25.0 |
| SK1d3 | 6.6 | 184 | 90.46 | 1.3 | 0.04 | 39.5 | 7.17 | 25.3 |
| LK1d3 | 5.4 | 208 | 19.47 | 3.06 | 0.1 | 8.5 | 7.52 | 25.2 |
| CK1d3 | 4.8 | 80 | 81.3 | 1.89 | 0.13 | 35.5 | 8.1 | 25.1 |
| Average,<br>d3 | 6 | 199.1 | 83.9 | 2.8 | 0.1 | 25.8 | 8.6 | 25.2 |

**Legend:** SG1-Sandy loam soil with *Aspilia pruliseta* vegetation at Gakurungu site; LG1-Silt loam soil with *Aspilia pruliseta* vegetation at Gakurungu site; CG1-Silty clay soil with *Aspilia pruliseta* vegetation at Gakurungu site; ST1- Sandy loam soil with *Aspilia pruliseta* vegetation at Tunyai site; LT1-Silt loam soil with *Aspilia pruliseta* vegetation at Tunyai site; CT1- Silty clay soil with *Aspilia pruliseta* vegetation at Tunyai site; SK1- Sandy loam soil with *Aspilia pruliseta* vegetation at Kanyuambora site; LK1- Silt loam soil with *Aspilia pruliseta* vegetation at Kanyuambora site; CK1- Silty clay soil with *Aspilia pruliseta* vegetation at Kanyuambora site; d<sub>3</sub>-Depth three (41-60 cm); MF-Mycorrhiza fungi; P<sub>2</sub>O<sub>5</sub>-Soil phosphates; SM-Soil moisture; N-Nitrogen; P-Phosphorus; OM-Soil organic matter

### Sample collection

Soils were sampled from Gakũrũngũ, Tunyai and Kanyuambora field sites using a standard soil auger (SOD-GP Dormer sampling equipment). A reconnaissance survey was initially carried out in which natural seed-bearing *Aspilia pruliseta* plots were mapped out. Besides the seed bearing vegetation of interest, the selected areas of the survey for each of the three sites had to have three soil textural types (sandy loam, silt loam and silty clay). In each site, a quadrant measuring one metre by one metre was thrown at random in each sub-site (a sub-site consisted of an area within the site with one soil textural type). In case the quadrant contained more than one *Aspilia pruliseta* plant, the one closest to the centre of the quadrant was chosen for collection of rhizosphere soil fungal spores. The quadrant was thrown five times in each sub-site and soil was sampled at depth_1_, 0-20 cm; depth_2_, 21-40 cm, depth_3_, 41-60 cm using a soil auger with a scooping capacity of 40cm^3^ of soil. The sampled soil was put together for each rhizosphere depth from each sub-site and homogenously mixed. 100 g of the mixture was put into khaki paper bags for soil and root DNA analysis in the laboratory.

### Soil and rootlets total DNA extraction

10 g of composite soil from the rhizosphere of *Aspilia pruliseta* comprising of the plant’s rootlets was weighed using an electronic balance. The weighed soil was put into 100 ml beaker and about 50 ml tap water was added. The mixture was placed and mixed on an electronic stirrer overnight. After 12 hours the mixture was washed several times by passing it through a 710 μ sieve placed on top of a 45 μ sieve (14). The 710 μ sieve collected the roots and course debris while the 45 μ prevented the spores from passing through. The roots and course debris from the 710 μ sieve were put into a mortar and air-dried in a hood while the process of sieving continued by collecting sieved water and soil mixture in a 1-litre cylinder (14). The washing and decanting process was done several times until near-clear water was obtained. This was followed by filling the centrifuge tubes with the sieved content. Centrifugation was done for 5 minutes at 1500 revolutions per minute (rpm) and the filtrate was poured off while the supernatant remained at the bottom of the tube. 48 % sucrose solution was added to the supernatant at equal volumes (50ml) and centrifuged for 1 minute at 1500 (rpm) (14). The filtrate was collected on the 45 μ sieve while the supernatant was disposed off. The filtrate was then washed with slowly flowing tap water to wash off the sucrose. The washed content was then collected in a 50ml plastic cylinder and the contents poured into a filter paper. Using a fine pair of forceps, the contents were picked and transferred to eppendorf tubes. The dried plant roots in the mortar were crushed into a fine powder using a pestle and the contents added to fungal spore cells in the eppendorf (20). The content in the eppendorf was re-suspended in 100 μl of solution A {100mM Tris-HCL (pH 8.0), 100Mm EDTA (pH 8.0); added to 5 μl of lysozyme (from a 20mg/ml solution) and incubated at 37^0^C for 30 minutes in a water bath}. 400 μl of lysis buffer (solution B) comprising 400 mM Tris-HCL (pH 8.0), 60 mM EDTA (pH 8.0), 150 mM NaCl, 1% sodium dodycyl sulfate and the tube was left at room temperature for 10 minutes. 10 μl of Proteinase K (20mg/ml) was mixed gently and incubated at 65^0^C for 1 hour in a water bath. An equal volume of chloroform/ isoamyl alcohol was added and centrifuged at 13200 rpm for 5 minutes at 4^0^C. The supernatant was transferred to new tubes. In the new tubes, 150 μl of sodium acetate (pH 5.2) and an equal volume of isopropanol alcohol was added accordingly. The tubes were briefly mixed through inversion. The mixture was then incubated at –20^0^C overnight. The tubes were then spun at 13200 rpm for 30 minutes and the supernatant was discarded. The resultant DNA pellets were washed in 300 μl of 70% ethanol. The pellets were then spun at 10000 rpm for 1 minute and the supernatant discarded. The resultant DNA pellet was air dried in the hood and dissolved in 50 ml of 13 Tris-EDTA. Genomic DNA (5–15 ng) in 10 μl of ddH2O was used for RAPD amplification using 1.5% agarose gels and images obtained confirming presence of DNA (20). 6 samples (3 samples each from depth_1_, 0-20 cm; depth_2_, 21-40 cm and depth_3_, 41-60 cm) were dried using LABCONCO machine. About 30 μl of the confirmed DNA was shipped to mrdnalabs (USA) for next generation sequencing with the primers as diversity assay bTEFAP® average inhouse ITSwanda. Illumina was used as the sequencing technology method.

### Amplicon library preparation and sequencing

Amplification of the ITS region on PCR was done using ITS1 (TCCGTAGGTGAACCTGCGG) and TS4 (TCCTCCGCTTATTGATATGC) primers with barcode according to (21). Amplification proceeded in a 30 cycle PCR using the HotStarTaq Plus Master Mix Kit (Qiagen, USA) with initial heating at 94°C for 3 min, followed by 28 cycles of denaturation at 94°C for 30 s, annealing at 53°C for 40 s and extension at 72°C for 1 min, after which a final elongation step at 72°C for 5 min was performed. Polymerase chain reaction (PCR) products were visualized on 2% agarose gel to determine the success of amplification and the relative intensity of bands. Multiple samples were pooled together in equal proportions based on their DNA concentrations. Pooled samples were purified using calibrated Ampure XP beads (Agencourt Bioscience Corporation, MA, USA). The pooled and purified PCR product was used to prepare DNA library according to Illumina sequencing protocol (22). Sequencing was performed at Molecular Research DNA (www.mrdnalab.com, Shallowater, TX, USA) on a MiSeq platform following the guidelines of the manufacturer.

### Sequence analysis, taxonomic classification and data Submission

Sequences obtained from the Illumina sequencing platform were depleted of barcodes and primers using a proprietary pipeline (www.mrdnalab.com, MR DNA, Shallowater, TX) developed at the service provider’s laboratory. Low quality sequences were identified by denoising and filtered out of the dataset according to (23). Sequences which were < 200 base pairs after phred20-based quality trimming, sequences with ambiguous base calls, and those with homopolymer runs exceeding 6bp were removed. Sequences were analyzed by a script optimized for high-throughput data to identify potential chimeras in the sequence files, and all definite chimeras were depleted as previously described (24). De novo OTU clustering was done with standard UCLUST method using the default settings as implemented in QIIME pipeline Version 1.8.0 at 97% similarity level (25). Taxonomy was assigned to each OTU using BLASTn against SILVA SSU Reference 119 database at default e-value threshold of 0.001 in QIIME (26).

### Data analysis

Shannon, Simpson and Evenness diversity indices were used for each sample and were calculated using vegan package version 1.16-32 in R software version 4.0.2 (27). Community and Environmental distances were compared using Analysis of similarity (ANOSIM) test, based upon Bray-Curtis distance measurements with 999 permutations. Significance was determined at 95% confidence interval (p<0.05). Calculation of Bray-Curtis dissimilarities between datasets and hierarchical clustering were carried out using the R programming language (27) and the Vegan package (28). To support OTU-based analysis, taxonomic groups were derived from the number of reads assigned to each taxon at all ranks from domain to genus using the taxa_summary.txt output from QIIME pipeline Version 1.8.0. Obtained sequences were submitted to NCBI Sequence Read Archive with SRP# Study accessions: SRP061806.

## Results

Soil pH for the three studied depths was slightly acidic with a range from 5.9-6.1 (Table 1a, 1b & 1c). The middle depth (21-40 cm, Table 1b) was more acidic with a pH of 5.9 compared to 6.1 and 6.0 in the first and third soil depth respectively. There was more organic matter content in depth two at 9.1% compared to 8.8% and 8.6% for depth one and depth three respectively. Soil temperatures declined with increasing rhizosphere depth from 25.5^0^C in depth one to 25.4^0^C and 25.2^0^C in depth two and three respectively. Mycorrhiza fungi (MF) spore counts along the rhizosphere of *Aspilia pruliseta* plant had an inverse relationship to soil depth with the top soil, depth one having 624 spores per 10g of the sample soil tested compared to 325.3 spores in depth two and 199.1 spores in depth three.

### Sequence data

Raw data consisted of *Aspilia pruliseta* rhizosphere soil samples taken in depth one (0-20 cm), depth two (21-40 cm) and depth three (41-60 cm) consisting of 271582 sequences of which 175622 were retained after removing sequences with different tags at each end for quality filtering and denoising. After removing singletons, chimeric sequences and OTUs of non-fungal organisms (<200 base pairs after phred20-based quality trimming, sequences with ambiguous base calls, and those with homopolymer runs exceeding 6 bp), a total of 373 OTUs were recovered at 3% genetic distance. 330 OTUs were of fungal origin and were further analysed.

### Diversity and Composition of fungal communities in the rhizosphere of *Aspilia pruliseta*

Based on BLASTn searches in SILVA SSU Reference 119 database, 323 fungal OTUs were identified, most of which had their best matches against accessions in SILVA database. These 324 OTUs spanned 5 phyla namely; Glomeromycota, Basidiomycota, Chytridiomycota, Ascomycota and unspecified phylum of fungi.

MC2_a_ that consisted of rhizosphere soil depth of 21-40 cm had the highest overall number of OTUs (283 OTUs) while MC1a (0-20 cm) and MC3_a_ (41-60 cm) had 262 and 265 overall OTUs respectively. 160 OTUs were shared among all sample types (Figure 1).

**Figure 1.**
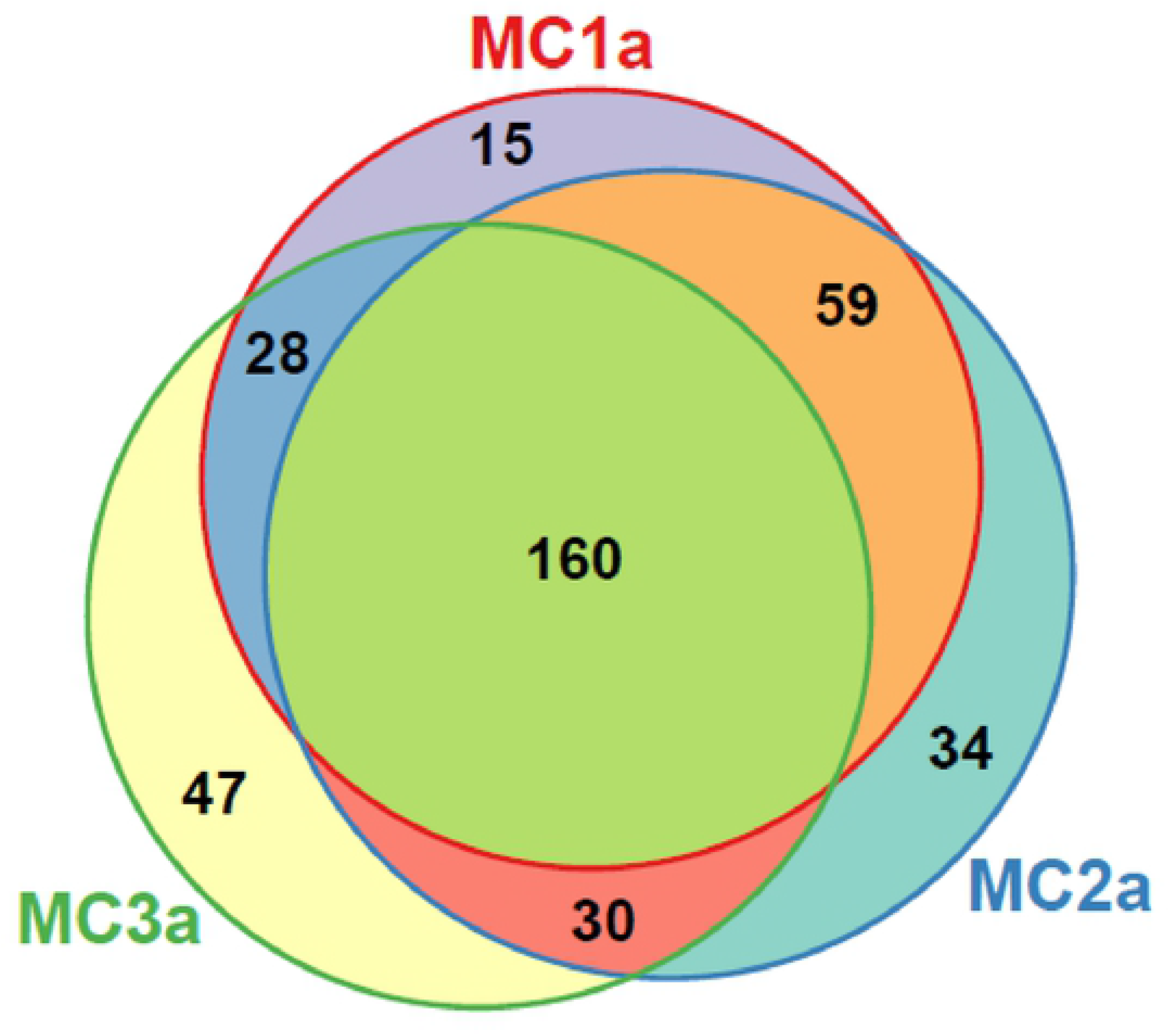
Venn diagram showing the distribution of unique and shared OTUs within various sample types in the three sampling sites. The number of OTUs in each rhizosphere depth is indicated in the respective circle.

Fungal OTUs were distributed among the phyla as follows; Glomeromycota (90.7%), Basidiomycota (3.7%), Ascomycota (3.4%), Chytridiomycota (1.5%), and unspecified phylum fungi (0.7%). Fungal phylum Glomeromycota was more abundant in rhizosphere depth one (0-20 cm) with 232 OTUs compared to depth two (21-40 cm) and depth three (41-60 cm) which had 229 and 213 OTUs respectively. This phylum was represented by most genera as shown in figure 2. The phylum Ascomycota had inverse OTU numbers to soil depth. At soil depth 0-20 cm, the phylum had 2 OTUs. The phylum had 5 and 9 OTUs at soil depth 21-40 cm and 41-60 cm respectively. Chytridiomycota and Basidiomycota fungal phylum had similar characteristics to Ascomycota and tended to inhabit the lower rhizosphere echelons. At 0-20 cm soil depth, OTUs were affiliated to the genus *Glomus* with a relative abundance of 85.3%, *Septoglomus* with 5.5% and *Paraglomus* with 4.9% whereas at 21-40 cm, the dominant genus was *Glomus* with relative abundance of 78.3% and *Rhizophagus* with a relative abundance of 15.8% while at soil depth 41-60 cm the dominant genus was *Glomus* with a relative abundance of 50.9% and *Septoglomus* with a relative abundance of 38.6% (Figure 2). The dominant species in the rhizosphere were *Glomus sp* and *Paraglomus laccatum.* The soil sample collected at soil depth 0-20 cm (MC1a) was found to harbor a higher diversity of fungi with low species richness as shown in Figure 2.

**Figure 2.**
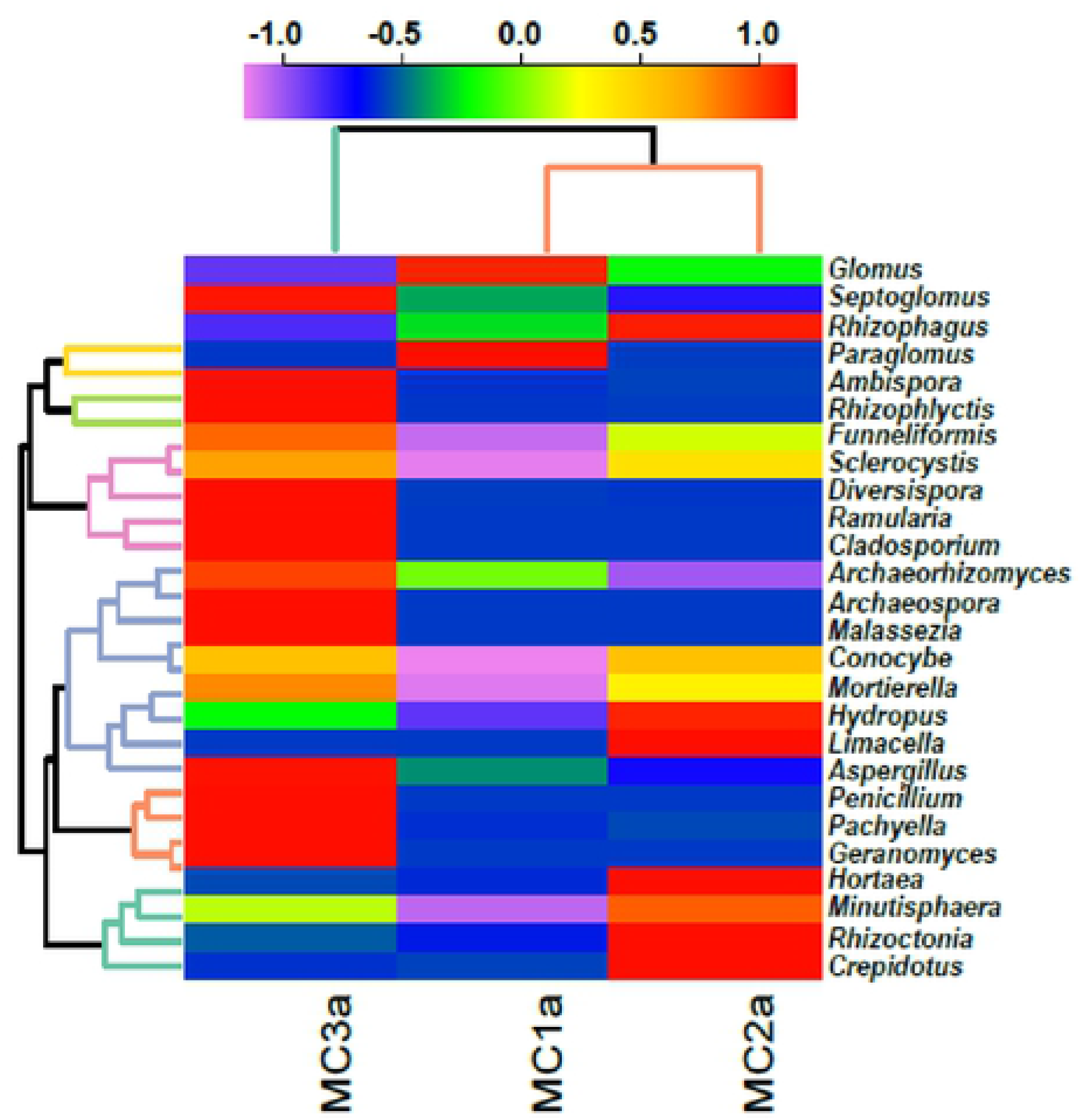
Heat map showing relative abundance of the most predominant fungal genera in various samples collected from *Aspilia pruliseta* rhizosphere.

Hierarchical clustering between samples collected from the rhizosphere of *Aspilia pruliseta* revealed samples from the second and third studied soil levels (21-60 cm) to be closer than from the sample in the first soil level, 0-20 cm (Figure 3). The dendogram shows relationships between the three samples collected.

**Figure 3.**
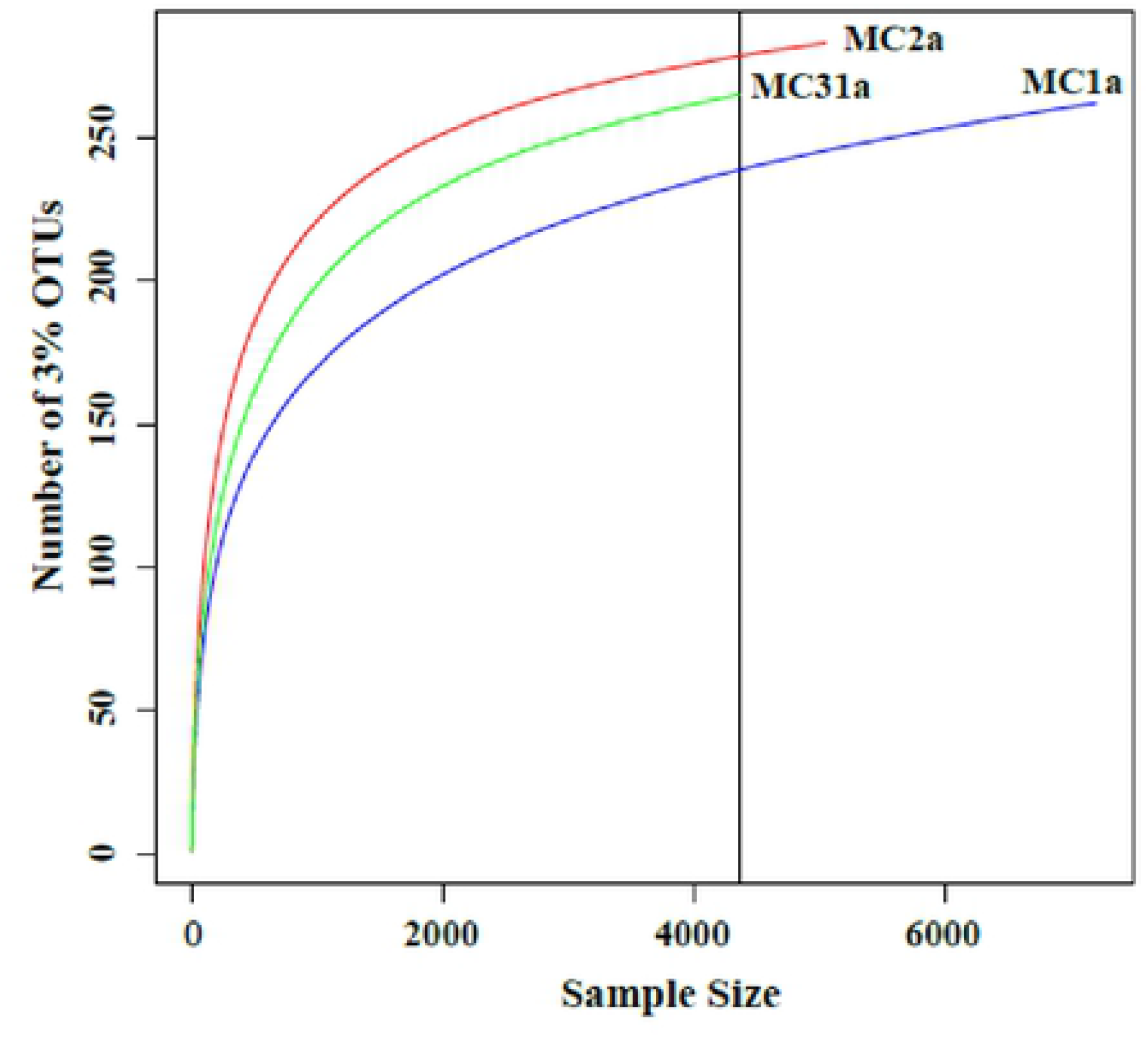
Relationships between the sample size sequenced and OTUs in the tested samples.

### Fungal richness and diversity indices

Richness (S) estimated the rhizosphere depth MC3a (41-60 cm) to be the richest site, constituting 62 taxa. Soil samples from the three sites had Evenness (J’) scores close to 0.1(0.0457 – 0.0978), hence showing evenness in their number of taxa members than the soil sample (41-60 cm). Simpson (1/D) also indicated the soil sample taken from depth 21-40 cm (MC2a) to harbor the most diverse taxa (12.808). The Shannon’s index (H’ = 2.48–3.32) indicated low variation in the level of diversity among the soil depth samples taken (Table 2).

**Table 2.** Diversity indices computed on all OTU-based fungal taxonomic units obtained from samples collected from the rhizosphere of *Aspilia pruliseta*

| Sample | Sequences after filtering | No. of OTUs | Richness (S) | Shannon (H) | Inverse Simpson (I/D) | Evenness (J) |
| --- | --- | --- | --- | --- | --- | --- |
| MC1a | 72,093 | 283 | 42 | 2.484 | 4.836 | 0.0457 |
| MC2a | 50,539 | 262 | 58 | 3.321 | 12.808 | 0.0978 |
| MC3a | 43,596 | 265 | 62 | 2.936 | 7.294 | 0.0711 |
| **Totals** | **166,228** | **323** | **162** |  |  |  |

Analysis of similarity and distance based redundancy analysis at class (Figure 4) level showed connectivity of distance matrix with threshold dissimilarity of 1 indicating that data of the three samples are connected ([1] 1 1 1), hence there were no significant differences in community structure in the samples at 95% level of confidence (P value=0.05).

**Figure 4.**
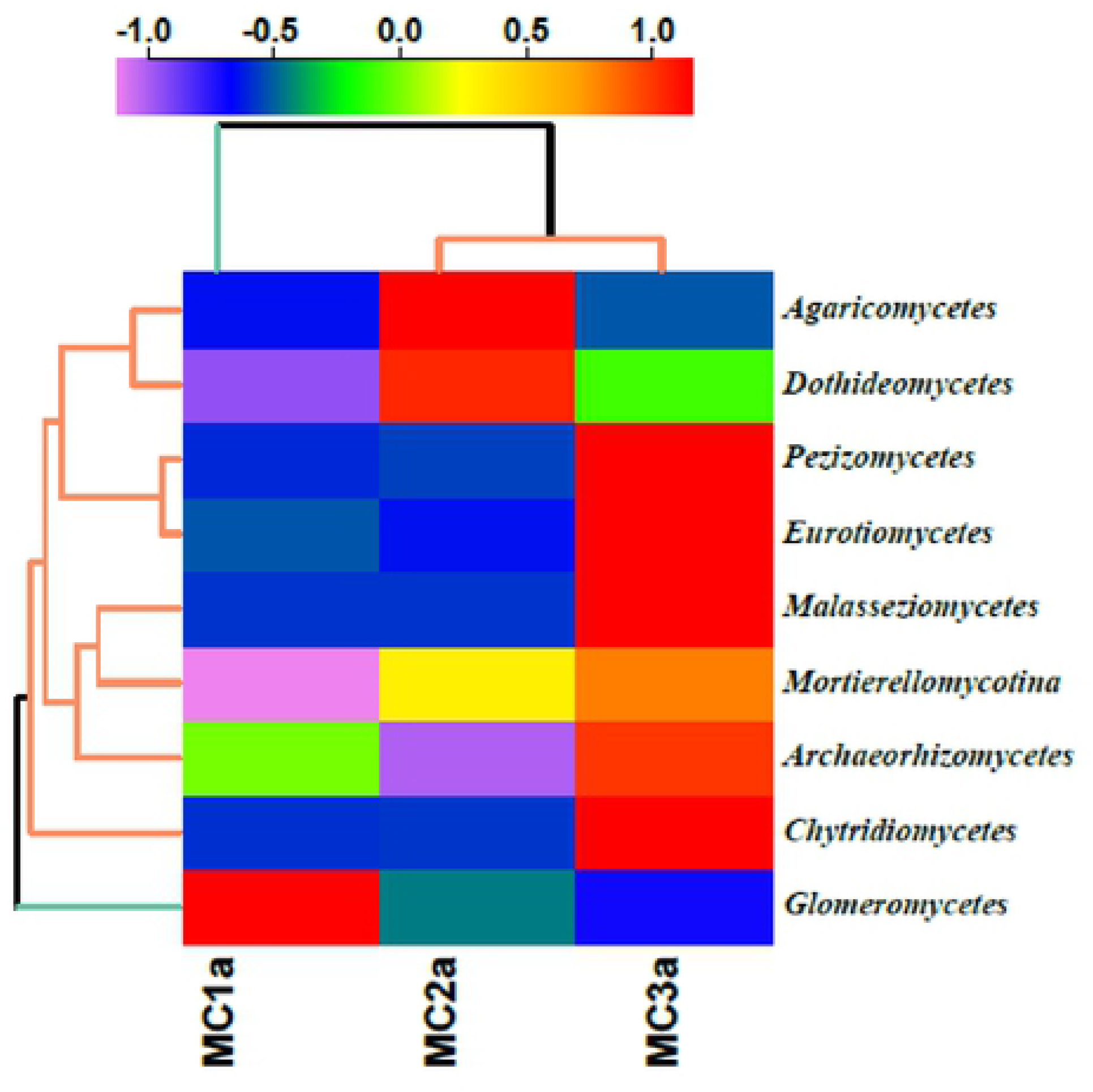
Hierarchical clustering of DNA samples collected from the studied soil rhizosphere depth. Class level was chosen to be used in hierarchical clustering to assess the relationships between samples and taxa.

## Discussion

Soil microbial community is responsible for most nutrient transformations in soil, regenerating minerals that limit plant productivity (29). Soil pH strongly influences fungal biomass composition (30). In this experiment, moderately acidic and sandy loam textured soils tended to favour proliferation of rhizosphere fungal growth (Table 1a, b &c). The level of soil organic matter was higher in the second rhizosphere layer (21-40 cm) but fungal microbe population was not correspondingly high agreeing with the principal findings of (31) that soil has diverse elements that contribute to its productivity and the proper balance between those elements is what actually matters.

The high sensitivity of Illumina sequencing enabled detection of rare species, thus providing more detailed information on fungal diversity in the rhizosphere of *Aspilia pruliseta* plant. The phylum, *Glomeromycota* was more frequently identified in the plant’s rhizosphere than those of *Basidiomycota* and *Ascomycota* whereas members of *Chytridiomycota* were represented on a smaller proportion of the rhizosphere fungal communities. The presence of unidentified fungal phylum indicate that new and potentially useful fungal communities do exist. Results from most rhizosphere mycological research findings indicate heavy presence of *Ascomycota* and *Basidiomycota* phyla (11) (12) and (13) from cultivated crops. From this research, there is a clear departure on the hierarchical fungal composition of the wild semi-arid shrub (*Aspilia pruliseta*) that could prove beneficial to follower-cultivated crops.

## Conclusion

This study presented fungal diversity analysis of rhizosphere soil samples collected from *Aspilia pruliseta* in the semi-arid eastern Kenya using Illumina Sequencing Technology. The results revealed heavy presence of phosphate solubilizing fungi suggesting the usefulness of the shrub for use in improving fallows. Optimal physico-chemical properties for AMF proliferation include sandy loam soils at 0-20 cm rhizosphere depth, warm and moderately acidic. The phylum, *Glomeromycota* dominated the plant’s rhizosphere depth.

## ACKNOWLEDGEMENT

Authors acknowledge funding in carrying out this research by National Research Fund (NRF) from the Kenya government.

## REFERENCE

1. Yuvaraj M, Ramasamy M. Role of Fungi in Agriculture. In 2020.

2. Anwar MS, Siddique MT, Verma A, Rao YR, Nailwal T, Ansari MW, et al. Multitrait plant growth promoting (PGP) rhizobacterial isolates from Brassica juncea rhizosphere: Keratin degradation and growth promotion. Commun Integr Biol. 2014;7(1):37–41.

3. Shukla A, Kumar A, Jha A, Ajit, Rao DVKN. Phosphorus threshold for arbuscular mycorrhizal colonization of crops and tree seedlings. Biol Fertil Soils. 2012;48(1):109–16.

4. Shukla A, Kumar A, Jha A, Salunkhe O, Vyas D. Soil moisture levels affect mycorrhization during early stages of development of agroforestry plants. Biol Fertil Soils. 2013;49(5):545–54.

5. Van Der Heijden MGA, Klironomos JN, Ursic M, Moutoglis P, Streitwolf-Engel R, Boller T, et al. Mycorrhizal fungal diversity determines plant biodiversity, ecosystem variability and productivity. Nature. 1998;396(6706):69–72.

6. Sreenivasa MN, Bagyaraj DJ. Use of pesticides for mass production of vesicular-arbuscular mycorrhizal inoculum. Plant Soil. 1989;119(1):127–32.

7. Brundrett MC. Understanding the Roles of Multifunctional Mycorrhizal and Endophytic Fungi. Microb Root Endophytes. 2007;9:281–98.

8. Schreiner RP, Mihara KL, McDaniel H, Bethlenfalvay GJ. Mycorrhizal fungi influence plant and soil functions and interactions. Plant Soil. 1997;188(2):199–209.

9. Jarošík V, Kováčiková E, Maslowská H. The influence of planting location, plant growth stage and cultivars on microflora of winter wheat roots. Microbiol Res. 1996;151(2):177–82.

10. Smit E, Leeflang P, Glandorf B, Van Elsas JD, Wernars K. Analysis of fungal diversity in the wheat rhizosphere by sequencing of cloned PCR-amplified genes encoding 18S rRNA and temperature gradient gel electrophoresis. Appl Environ Microbiol. 1999;65(6):2614–21.

11. Zimudzi J, Waals JE Van Der, Coutinho TA, Cowan DA, Valverde A. Temporal shifts of fungal communities in the rhizosphere and on tubers in potato fields Josephine. Fungal Biol [Internet]. 2018; Available from: https://doi.org/10.1016/j.funbio.2018.05.008

12. Jie W, Lin J, Guo N, Cai B, Yan X. Community composition of rhizosphere fungi as affected by Funneliformis mosseae in soybean continuous cropping soil during seedling period. 2019;79(September):356–65.

13. Floc J, Hamel C, Harker KN, St-arnaud M. Fungal Communities of the Canola Rhizosphere : Keystone Species and Substantial Between-Year Variation of the Rhizosphere Microbiome. 2020;

14. Varma A (1998). BLMS science and business media. https://doi.org/10.1007/97.-3-642-60268-9. Varma, A. (1998). Biology Lab Manual. Springer science and business media. ht. In Springer science and business media; 1998. Available from: https://www2.dijon.inrae.fr/mychintec/Protocole/protoframe.html

15. Olsen SR, Cole C V, Watandbe F, Dean L. Estimation of Available Phosphorus in Soil by Extraction with sodium Bicarbonate. J Chem Inf Model. 1954;53(9):1689–99.

16. Johnson A. Methods of measuring Soil Moisture in the Field. Geol Surv Water-Supply Pap 1619-U [Internet]. 1962;112(January 2007):11–32. Available from: http://medcontent.metapress.com/index/A65RM03P4874243N.pdf

17. Bremner JM. Determination of nitrogen in soil by the Kjeldahl method. J Agric Sci. 1960;55(1):11–33.

18. Mildred SS. Colorimetric Determination of Phosphorus in Soils. Anal Chem. 1942;23(10):1496–7.

19. Schulte EE, Hoskins B. Recommended Soil Organic Matter Tests. Recomm Soil Test Proced Northeast United States. 2009;63–74.

20. Lee SB, Milgroom MG, Taylor JW. A rapid, high yield mini-prep method for isolation of total genomic DNA from fungi. Fungal Genet Rep. 1988;35(1):23.

21. White TJ, Bruns T, Lee S, Taylor J. Amplification and Direct Sequencing of Fungal Ribosomal Rna Genes for Phylogenetics. PCR Protoc. 1990;(January):315–22.

22. Yu K, Zhang T. Metagenomic and metatranscriptomic analysis of microbial community structure and gene expression of activated sludge. PLoS One. 2012;7(5).

23. Reeder J, Knight R. Rapid denoising of pyrosequencing amplicon data: exploiting the rank-abundance distribution. Nat Methods [Internet]. 2010;7(9):668–9. Available from: http://www.ncbi.nlm.nih.gov/pmc/articles/PMC2945879/

24. Gontcharova VEY, Sun1 Y, Wolcott2 RD, Dowd and SE. A Comparison of Bacterial Composition in Diabetic Ulcers and Contralateral Intact Skin. Open Microbiol J. 2010;4(1):8–19.

25. J Gregory Caporaso, Justin Kuczynski, Jesse Stombaugh, Kyle Bittinger, Frederic D Bushman, Elizabeth K Costello, Noah Fierer, Antonio Gonzalez Peña, Julia K Goodrich, Jeffrey Gordon, Gavin A Huttley, Scott T Kelley, Dan Knights5, Jeremy E Koenig RE. QIIME allows analysis of high-throughput community sequencing data. Nat Methods. 2010;7(5):1–12.

26. Quast C, Pruesse E, Yilmaz P, Gerken J, Schweer T, Yarza P, et al. The SILVA ribosomal RNA gene database project: Improved data processing and web-based tools. Nucleic Acids Res. 2013;41(D1):590–6.

27. Hothorn T, Everitt BS. - An Introduction to R. A Handb Stat Anal using R. 2020;2:32–55.

28. Oksanen J, Blanchet FG, Friendly M, Kindt R, Legendre P, Mcglinn D, et al. Package “vegan” Title Community Ecology Package. Community Ecol Packag [Internet]. 2019;2(9):1–297. Available from: https://cran.r-project.org/web/packages/vegan/vegan.pdf

29. Rousk J, Brookes PC, Bååth E. Contrasting soil pH effects on fungal and bacterial growth suggest functional redundancy in carbon mineralization. Appl Environ Microbiol. 2009;75(6):1589–96.

30. Fierer N, Jackson RB. The diversity and biogeography of soil bacterial communities. Proc Natl Acad Sci U S A. 2006;103(3):626–31.

31. Bhattarai B. Variation of Soil Microbial Population in Different Soil Horizons. J Microbiol Exp. 2015;2(2):75–8.

